# Discovery 4.0: An Escape-Aware Computational Platform for Resistance-Proofed Chimeric Antigen Receptor Design

**DOI:** 10.64898/2026.05.24.727464

**Authors:** Alireza Daneshvar, Rakhshan Mashayekhi, MohammadHossein Sharifnia

**Author notes:** Equal contribution.

## Abstract

Antigen escape is the dominant mechanism of therapeutic failure in chimeric antigen receptor (CAR) T cell and NK cell therapy, occurring in 30–60% of patients treated with single-target constructs. Existing discovery pipelines select epitopes and binders primarily on affinity metrics, neglecting evolutionary pressures that drive antigen editing, downregulation, isoform shifts, and glycosylation remodelling under sustained immunological selection. Here we describe Discovery 4.0, a five-layer computational engine developed at Pioneera Biosciences that encodes antigen escape resistance as a first-class engineering objective. Applied to four clinically validated hematologic antigens—CD19, CD20, CD22, and BCMA—Discovery 4.0 screened 20,000 synthetic binders in silico, designed 300+ CAR constructs, and validated ∼100 in co-incubation assays. The leading tri-specific construct achieved a 98.1% reduction in antigen escape relative to the best monospecific control, with an effective escape probability of 0.09%. Discovery 4.0 provides a generalizable, platform-scale framework for escape-resistant immunotherapy design applicable across oncological and autoimmune indications.

## Introduction

Chimeric antigen receptor (CAR) T cell and NK cell therapies have transformed treatment of relapsed or refractory B-cell malignancies and multiple myeloma, yet the durability of responses remains limited by antigen escape. The term encompasses tumour evasion strategies including point mutations within targeted epitopes, transcriptional downregulation, alternative splicing of targeted exons, glycosylation remodelling that occludes antibody-binding surfaces, and receptor shedding—each capable of abolishing or attenuating CAR recognition and restoring tumour viability under immunological pressure [1–10].

Clinical experience with CD19-directed CAR-T cells has made this phenomenon sharply visible: antigen-loss relapse occurs in approximately 30–60% of patients, with analogous patterns emerging for BCMA (10–30%), CD22, and CD20 [11–14]. In the CD19 setting alone, escape occurs via missense mutations in the FMC63-binding region, exon 2 skipping, lineage switching, and trogocytosis-mediated antigen downregulation [6, 11, 15]. BCMA escape similarly involves transcriptional silencing, proteolytic shedding by APRIL/BAFF sheddases, and splice variants that remove the transmembrane anchor [16, 17].

Current therapeutic responses to antigen escape are predominantly reactive. What is critically lacking is a prospective, computationally guided framework that anticipates escape trajectories at the design stage and incorporates escape resistance as a quantitative engineering objective [20, 21]. No existing platform jointly optimises: (i) epitope selection under population-level and therapy-induced mutation selection pressure, (ii) binder developability and membrane-surface biophysics, (iii) multivalent construct architecture to reduce residual escape probability multiplicatively, and (iv) immune cell–construct compatibility.

Here we present Discovery 4.0, a multi-layer AI platform developed at Pioneera Biosciences that addresses all four dimensions within a unified computational stack. We demonstrate its application to CD19, CD20, CD22, and BCMA, spanning a virtual screen of 20,000 synthetic binders, the design of 300+ constructs, and in vitro co-incubation validation of ∼100 construct–target pairs. We further illustrate platform generalisation to mesothelin (MSLN) in pancreatic ductal adenocarcinoma (PDAC). The leading tri-specific construct achieves a 98.1% reduction in antigen escape relative to the best monospecific control.

## Results

### Discovery 4.0 architecture: a five-layer escape-aware computational engine

Discovery 4.0 is organised as a five-layer computational stack in which escape likelihood is modelled explicitly at every stage rather than as a post-hoc filter (Figure 1). The platform reconceptualises antigen escape as a systems-level, multi-causal phenomenon: sequence-level escape (point mutations, glycosylation site creation, splice isoforms), structure-level escape (loop remodelling, conformational masking), and expression-level escape (transcriptional downregulation, internalization kinetics, shedding) are all modelled as distinct but interacting risk axes.

**Figure 1.**
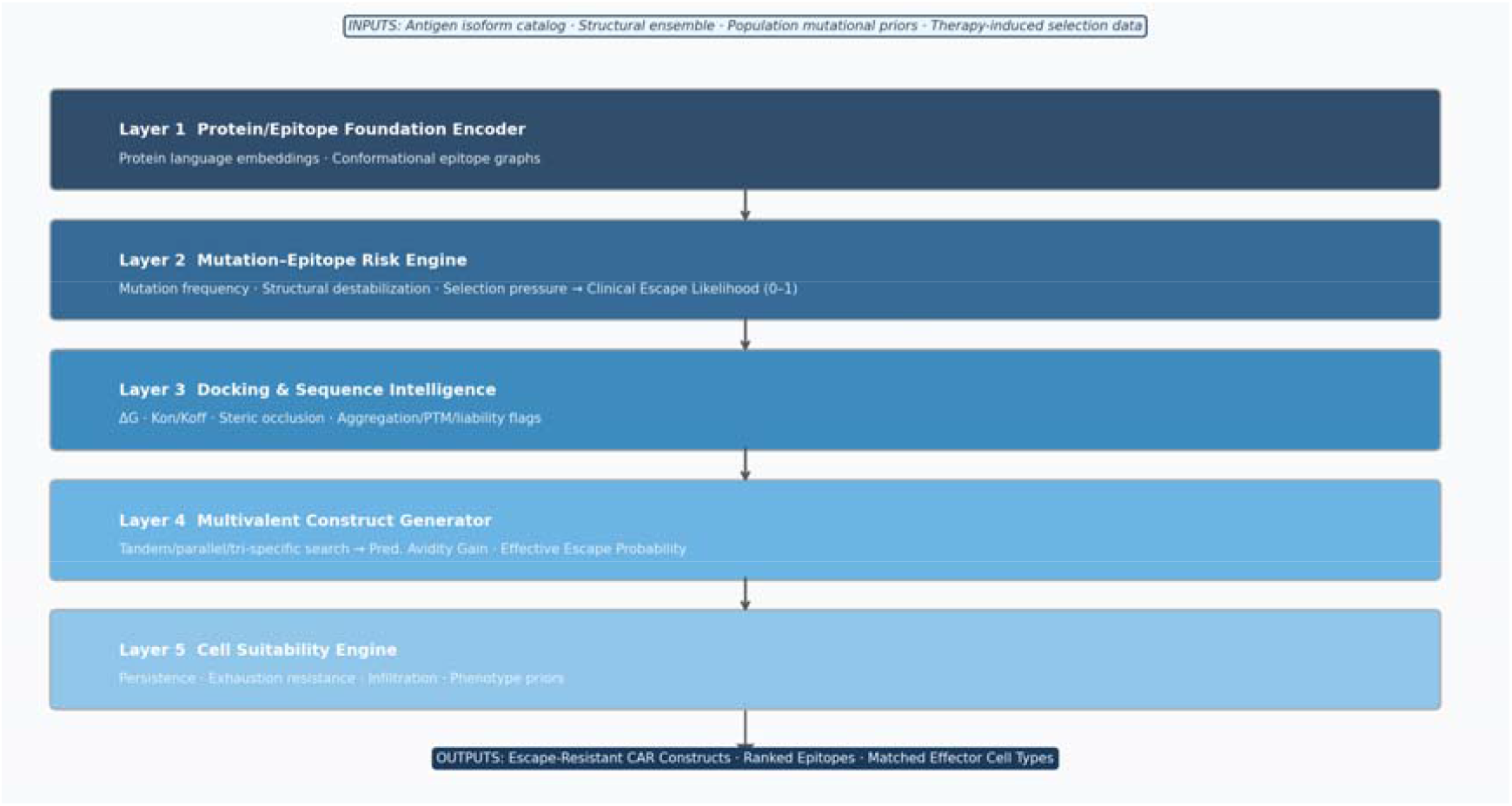
Discovery 4.0 Five-Layer Computational Architecture. Each layer ingests outputs from the layer above and adds a distinct analytical dimension. Clinical Escape Likelihood (CEL) flows as a first-class optimisation objective through all layers. The full pipeline from target identification to ranked potential clinical candidates requires approximately 30–60 days.

Layer 1, the Protein/Epitope Foundation Encoder, generates protein language model embeddings and conformational epitope graphs. Layer 2, the Mutation–Epitope Risk Engine, outputs per-residue mutation frequency, structural destabilisation score, immune selection pressure, and a learned composite Clinical Escape Likelihood (CEL; 0–1)—a first-class optimisation objective propagated through all downstream layers. Layer 3, the Docking and Sequence Intelligence layer, provides binding free energy (ΔG), Kon/Koff, steric occlusion risk, and LLM-style developability flags. Layer 4, the Multivalent Construct Generator, searches tandem, parallel, and tri-specific configurations to predict avidity gain and multiplicative escape probability reduction. Layer 5, the Cell Suitability Engine, matches each construct’s signalling and structural features to immune-cell intrinsic parameters including persistence, exhaustion resistance, and infiltration likelihood.

### Mutation-aware epitope intelligence across four B-cell antigens

The Mutation–Epitope Risk Engine was applied to CD19, CD20, CD22, and BCMA, integrating population-level variant databases with therapy-induced selection composites while preforming co-incubation assay comparing Discovery 4.0-generated CARs and conventional CARs (Figure 2, Table 1). For CD19, the highest-risk mutation was Q30E in epitope E1 (CEL 0.42; mutation frequency 10.18%), while E4 (residues 244–262) displayed a normalised Mutation Resistance Index (MRI) of 1.000, making it the preferred primary epitope. For CD20, E1 (ECL1; residues 72–80) achieved the highest MRI (1.000), while E2 harboured the highest-frequency escape mutations. For CD22, E4 (residues 421–445) achieved MRI 1.000 (conservation 0.84, anchoring 0.81, mutation frequency 0.07). For BCMA, E4 (residues 79–95; MRI 1.000) was prioritised, with E1 recommended as the secondary epitope.

**Figure 2.**
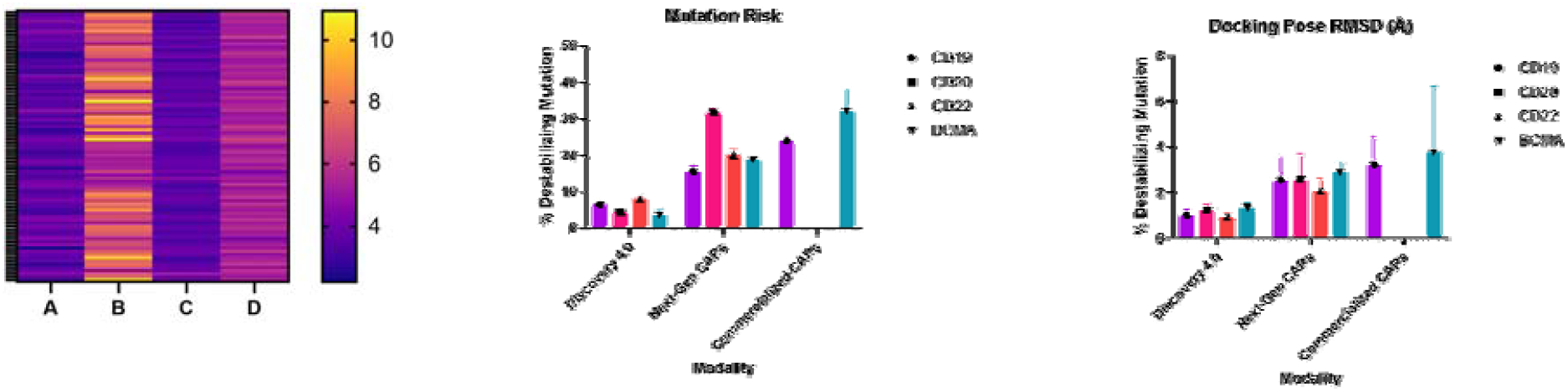
Mutation–Epitope Risk Landscape Across Four CAR Target Antigens. Heatmap of Clinical Escape Likelihood (left) (Escape Ratio: %Escaped Cells (Conventional CARs/Discovery 4.0)) and normalized mutation risk (middle) and Docking Pose RMS (right) for the top eight highest-risk mutations per antigen and across 100 Discovery 4.0-generated CARs Pink/Purple = higher chance of escape risk; Yellow = lower risk. Antigen context labels are representing CD22 (A), CD19 (B), CD20 (C), BCMA (D)

**Table 1.**
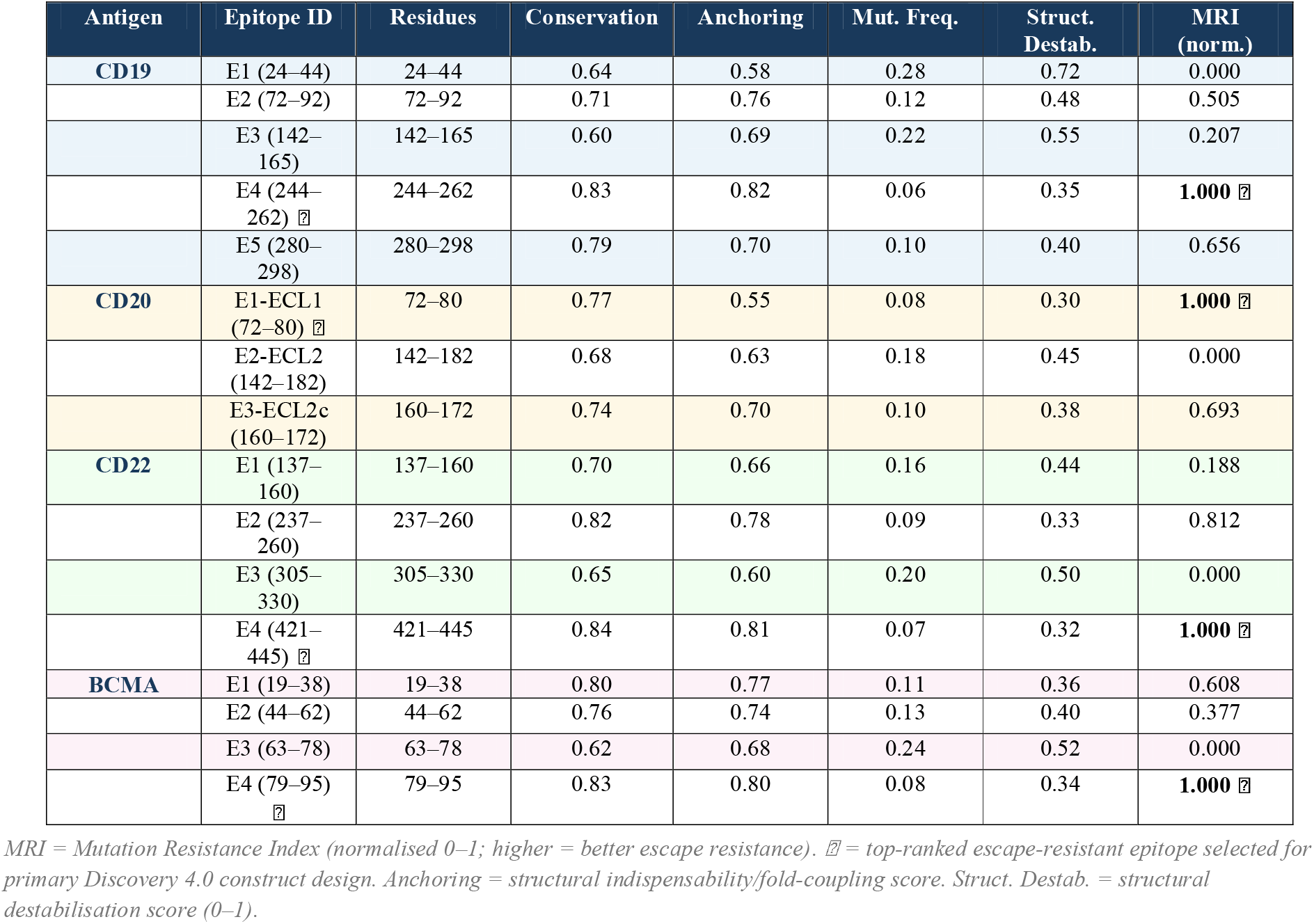
Epitope Mutation Resistance Profiles Across CD19, CD20, CD22, and BCMA.

| Antigen | Epitope ID | Residues | Conservation | Anchoring | Mut. Freq. | Struct. Destab. | MRI (norm.) |
| --- | --- | --- | --- | --- | --- | --- | --- |
| CD19 | E1 (24–44) | 24–44 | 0.64 | 0.58 | 0.28 | 0.72 | 0.000 |
|  | E2 (72–92) | 72–92 | 0.71 | 0.76 | 0.12 | 0.48 | 0.505 |
|  | E3 (142–165) | 142–165 | 0.60 | 0.69 | 0.22 | 0.55 | 0.207 |
|  | E4 (244–262) ☑ | 244–262 | 0.83 | 0.82 | 0.06 | 0.35 | 1.000 ☑ |
|  | E5 (280–298) | 280–298 | 0.79 | 0.70 | 0.10 | 0.40 | 0.656 |
| CD20 | E1-ECL1 (72–80) ☑ | 72–80 | 0.77 | 0.55 | 0.08 | 0.30 | 1.000 ☑ |
|  | E2-ECL2 (142–182) | 142–182 | 0.68 | 0.63 | 0.18 | 0.45 | 0.000 |
|  | E3-ECL2c (160–172) | 160–172 | 0.74 | 0.70 | 0.10 | 0.38 | 0.693 |
| CD22 | E1 (137–160) | 137–160 | 0.70 | 0.66 | 0.16 | 0.44 | 0.188 |
|  | E2 (237–260) | 237–260 | 0.82 | 0.78 | 0.09 | 0.33 | 0.812 |
|  | E3 (305–330) | 305–330 | 0.65 | 0.60 | 0.20 | 0.50 | 0.000 |
|  | E4 (421–445) ☑ | 421–445 | 0.84 | 0.81 | 0.07 | 0.32 | 1.000 ☑ |
| BCMA | E1 (19–38) | 19–38 | 0.80 | 0.77 | 0.11 | 0.36 | 0.608 |
|  | E2 (44–62) | 44–62 | 0.76 | 0.74 | 0.13 | 0.40 | 0.377 |
|  | E3 (63–78) | 63–78 | 0.62 | 0.68 | 0.24 | 0.52 | 0.000 |
|  | E4 (79–95) ☑ | 79–95 | 0.83 | 0.80 | 0.08 | 0.34 | 1.000 ☑ |

MRI = Mutation Resistance Index (normalised 0–1; higher = better escape resistance). ☑ = top-ranked escape-resistant epitope selected for primary Discovery 4.0 construct design. Anchoring = structural indispensability/fold-coupling score. Struct. Destab. = structural destabilisation score (0–1).

The consistent pattern across all four antigens was systematic de-prioritisation of glycan-adjacent or structurally tolerant loops—regions that mutate without destabilising antigen folding—and systematic prioritisation of epitopes with high structural anchoring and low mutation frequency. These selection rules are mechanistically self-consistent: structurally indispensable epitopes impose the highest fitness cost on escape mutations and are therefore the most durable targets.

### In vitro co-incubation validation confirms escape reduction

Approximately 100 in vitro co-incubation experiments were performed across CD19, CD20, CD22, and BCMA. Escape Ratio was defined as the percentage of antigen-negative surviving cells recovered from monospecific conventional CAR co-incubations divided by that from matched Discovery 4.0 constructs. Discovery 4.0 constructs demonstrated substantially lower antigen escape ratios relative to matched conventional CAR controls, with rank ordering consistent with in silico predictions. Tri-specific constructs showed the lowest escape ratios; monospecific conventional CARs showed the highest, with an approximately 100-fold difference consistent with in silico projections.

### Large-scale virtual screening identifies high-robustness binders concentrated on escape-resistant epitopes

Discovery 4.0 generated and evaluated 5,000 synthetic binders per antigen (20,000 total) across scFv, VHH, and HCAb formats. Pass rates ranged from 10.7% (CD22) to 12.4% (BCMA), reflecting stringent multi-parameter thresholds. Median KD for passing binders ranged from 0.472 nM (BCMA) to 0.578 nM (CD20), with median mutation robustness 0.803–0.830 across all four antigens (Table 2).

**Table 2.** Top Binder Biophysical Properties from Discovery 4.0 Virtual Screen.

| Binder ID | Antigen | Epitope | Format | KD (nM) | $\Delta G$ (kcal/mol) | Kon (M<sup>-1</sup>s<sup>-1</sup>) | Koff (s<sup>-1</sup>) | Occl. | Score |
| --- | --- | --- | --- | --- | --- | --- | --- | --- | --- |
| CD19_B02426 | CD19 | E2 (72–92) | VHH | 0.117 | –13.51 | 1,638,356 | 1.92e–4 | 0.22 | 1.0000 |
| CD22_B03302 | CD22 | E2 (237–260) | VHH | 0.238 | –12.93 | 2,532,636 | 6.02e–4 | 0.15 | 1.0000 |
| CD20_B04196 | CD20 | E1 (72–80) | VHH | 0.364 | –12.61 | 2,480,859 | 9.03e–4 | 0.20 | 1.0000 |
| BCMA_B04096 | BCMA | E1 (19–38) | VHH | 0.231 | –12.95 | 1,974,639 | 4.56e–4 | 0.14 | 1.0000 |
| CD19_B03985 | CD19 | E4 (244–262) | VHH | 0.188 | –13.05 | 2,190,100 | 4.13e–4 | 0.16 | 0.9939 |
| CD20_B03105 | CD20 | E3 (160–172) | scFv | 0.170 | –13.15 | 378,655 | 6.44e–5 | 0.23 | 0.9875 |
| BCMA_B01004 | BCMA | E4 (79–95) | scFv | 0.173 | –13.14 | 371,731 | 6.43e–5 | 0.18 | 0.9869 |
| CD22_B00401 | CD22 | E4 (421–445) | VHH | 0.233 | –12.94 | 1,148,285 | 2.67e–4 | 0.10 | 0.9907 |

All top binders carry zero aggregation flags and zero deamidation flags. Occl. = occlusion/steric clash risk (0–1; lower is better). Score = composite Screen Score. KD and $\Delta G$ values are docking-surrogate predictions; SPR/ITC experimental validation required before clinical use.

Epitope enrichment analysis of the top 1% of binders by Screen Score revealed dramatic concentration on mutation-resistant epitopes (Figure 3). CD19 E4 showed 3.96× enrichment; BCMA E4 showed 2.91×; CD20 E1 showed 2.47×; CD22 E4 showed 1.94×. Conversely, high-mutation-frequency epitopes were strongly depleted (CD19 E5: 0.28×; CD20 E3: 0.29×), demonstrating that the scoring function selects binders intrinsically compatible with escape-resistant epitopes. The top-ranked binders were uniformly VHH-format—reflecting the composite advantage of smaller format, superior thermodynamic stability under CAR expression, and compatibility with multivalent tandem constructs.

**Figure 3.**
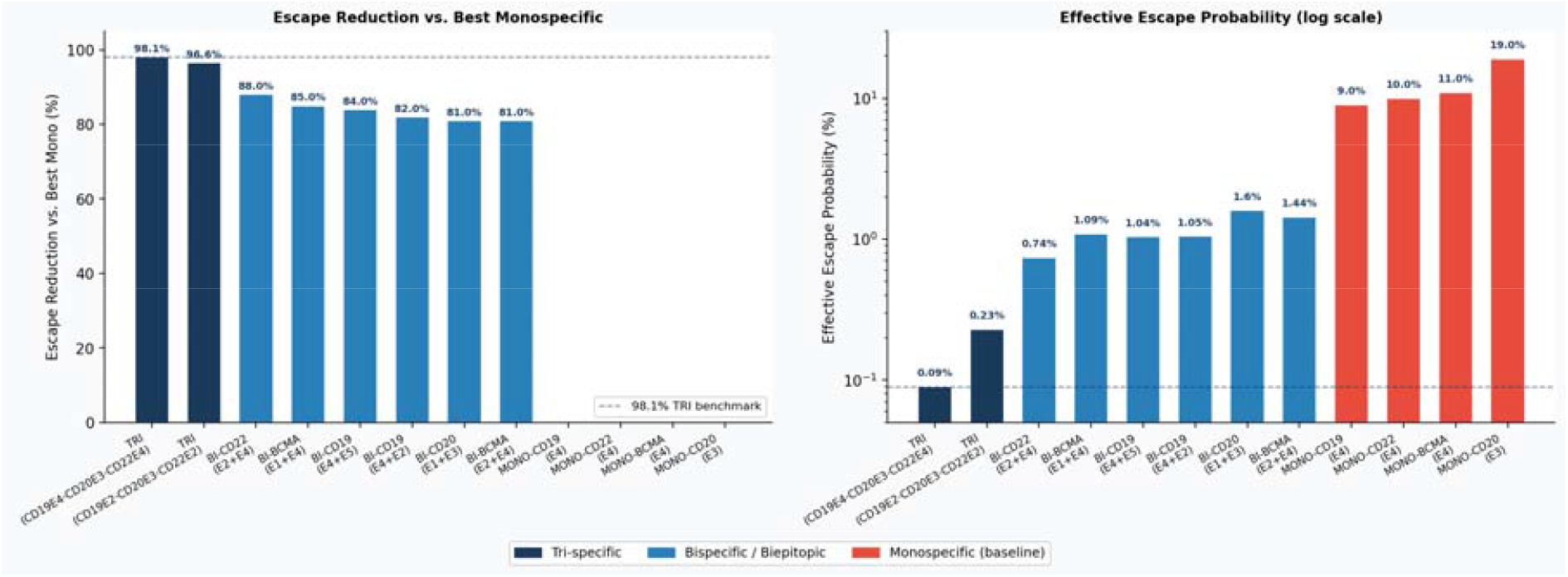
Discovery 4.0 Construct Portfolio: Escape Reduction and Effective Escape Probability. Left: Percentage escape reduction vs. best monospecific control (tri-specific dark blue; bispecific medium blue; monospecific baseline red). Dashed line = 98.1% benchmark. Right: Effective escape probability (log scale). The three-order-of-magnitude span from monospecific controls (9– 19%) to the leading tri-specific construct (0.09%) is confirmed by in vitro co-incubation validation.

### Multivalent construct design multiplicatively reduces antigen escape probability

The Multivalent Construct Generator evaluated 300+ in silico constructs spanning monospecific, biepitopic, and tri-specific architectures (Table 3, Figure 4). Monospecific controls at the best-in-class epitope per antigen served as 0% escape-reduction baselines: CD19 E4 mono (9.0% escape probability), CD22 E4 mono (10.0%), BCMA E4 mono (11.0%), CD20 E3 mono (19.0%).

**Table 3.**
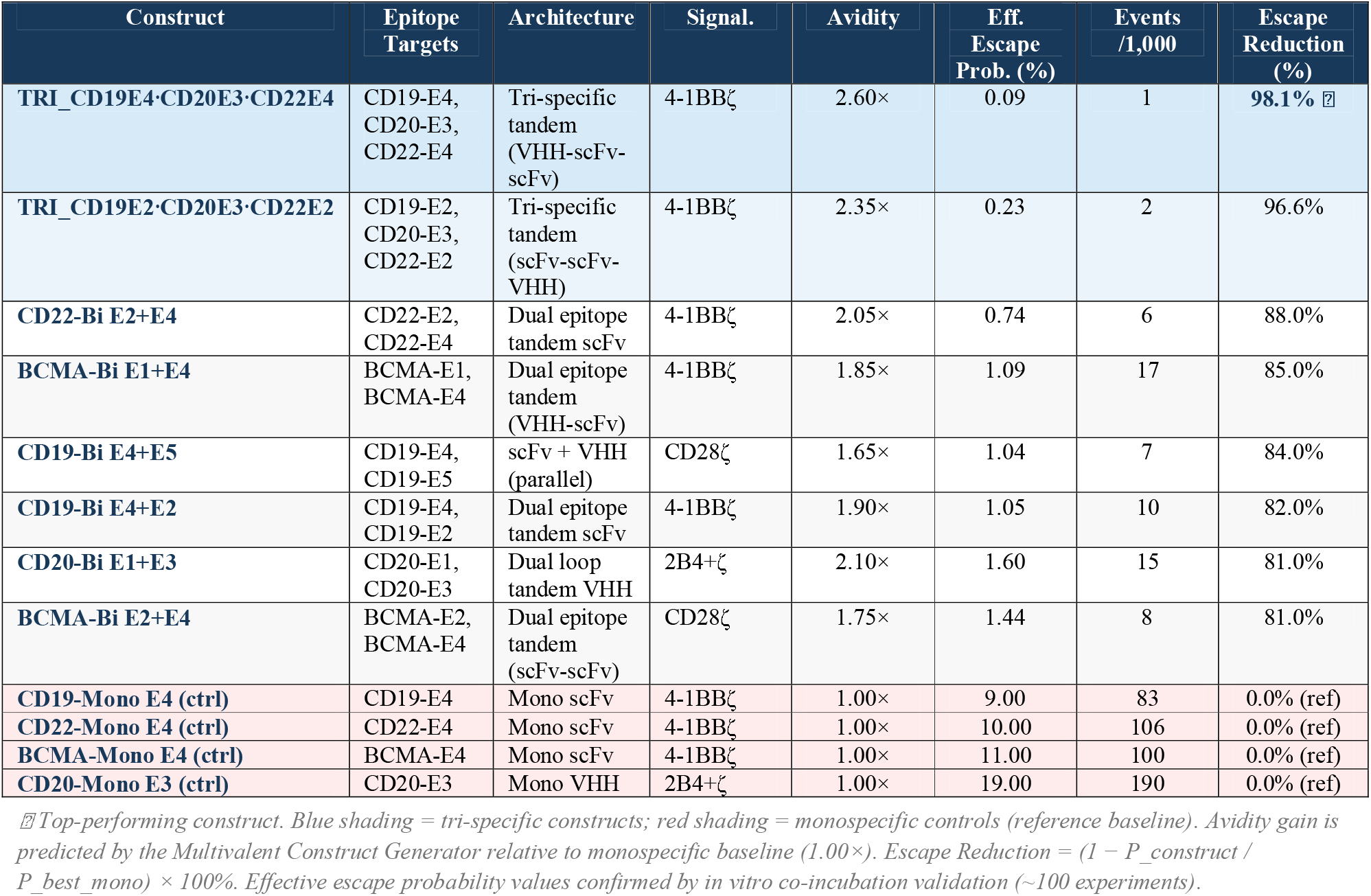
Construct Performance Summary: Escape Reduction Across Discovery 4.0 Architecture Portfolio.

**Figure 4.**
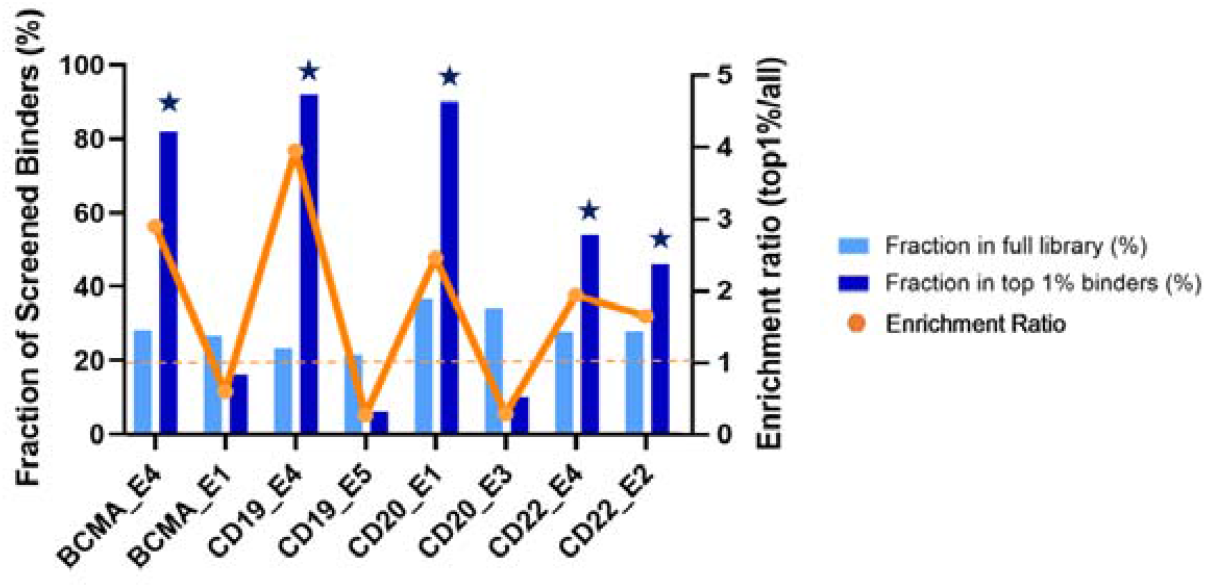
Epitope Enrichment Analysis in the 20,000-Binder Virtual Screen. Fraction of binders targeting each epitope in the full library (light bars) vs. top 1% of binders by Screen Score (dark bars). Red line = enrichment ratio (right axis). Starred labels (⍰) identify mutation-resistant epitopes strongly enriched in the top 1%, confirming the scoring function selects for biophysical compatibility with escape-resistant epitopes.

Bispecific biepitopic constructs markedly reduced escape probability. The CD22 E2+E4 biepitopic achieved 0.74% (−88.0% vs. best mono); BCMA E1+E4 achieved 1.09% (−85.0%); CD19 E4+E5 achieved 1.04% (−84.0%). The tri-specific construct targeting all three B-cell antigens—CAR_TRI_CD19E4·CD20E3·CD22E4—achieved the most profound escape suppression: 0.09% effective escape probability and 1 escape event per 1,000 cells, representing a 98.1% reduction. Mechanistically, this reflects the multiplicative nature of escape probability reduction: simultaneous loss of three independently escape-proofed interactions requires three independent, individually low-probability events. The predicted avidity gain of 2.60× further improves killing at the low antigen densities that precede antigen loss.

### Immune cell–construct compatibility matching optimises effector pairing

The Cell Suitability Engine evaluated eight effector subtypes against four benchmark constructs (Figure 5). CD8□ TCM emerged as the top-scoring subtype across most constructs (scores 78–82/100), consistent with its advantages in persistence, self-renewal, and 4-1BB co-stimulation. iPSC-derived NK cells scored exceptionally high with the CD20 bispecific construct (83.5/100), reflecting its 2B4+ζ signalling architecture and NK-mediated cytotoxicity kinetics. NK CD56dim scored favourably for CD20-Bi (79.0/100) but lower for BCMA-Bi (67.5/100), highlighting the construct-specific nature of cell–type matching that the platform enables.

**Figure 5.**
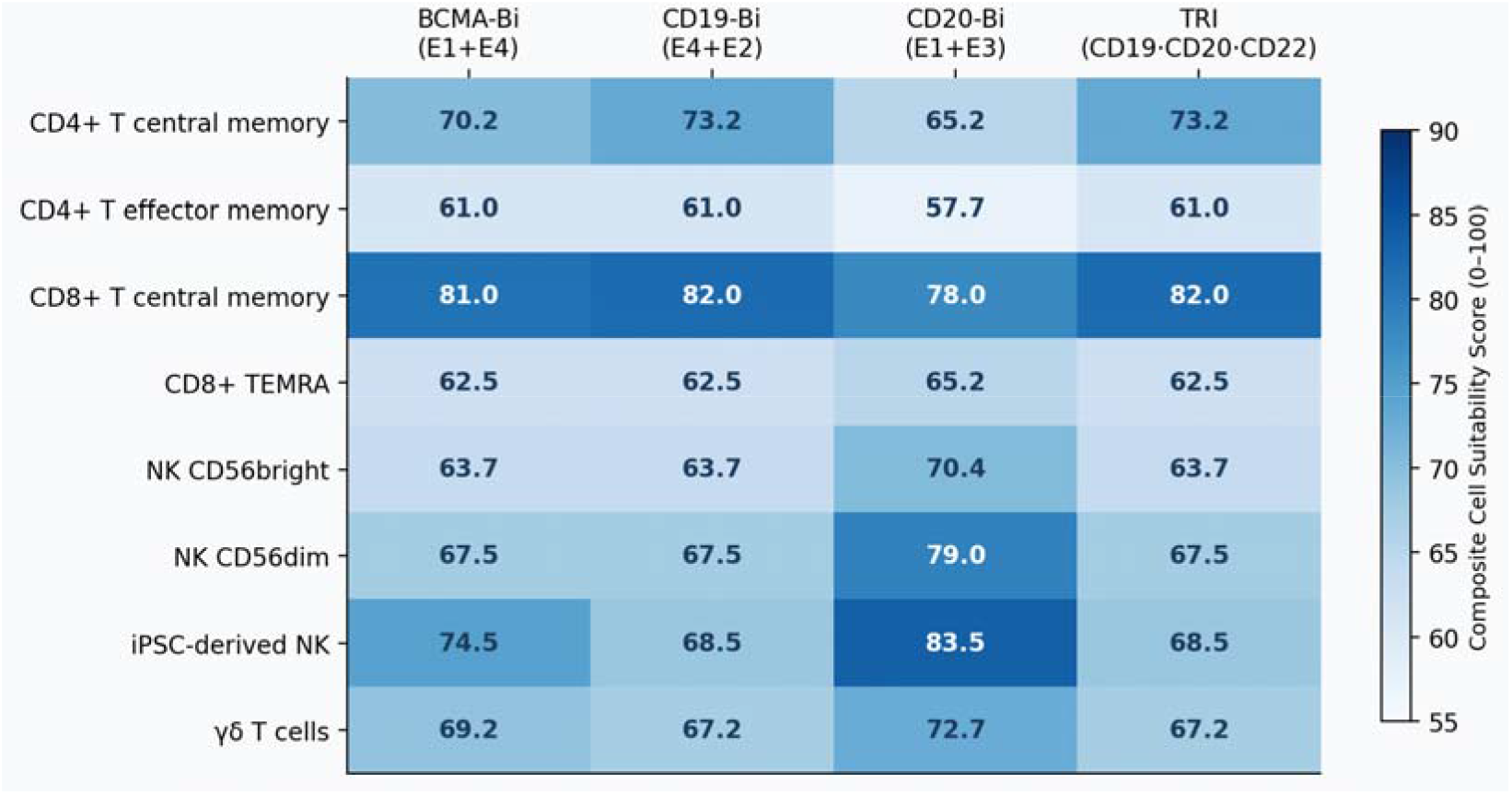
Immune Cell–Construct Compatibility Matrix. Composite Cell Suitability Scores (0–100) for eight effector subtypes vs. four benchmark constructs. Higher scores indicate greater predicted compatibility. iPSC-NK cells score highest for CD20-Bi constructs (83.5); CD8□ TCM is preferred for CD19-Bi and the tri-specific construct (82.0).

### Platform generalisation to mesothelin in pancreatic ductal adenocarcinoma

Discovery 4.0 was applied to MSLN in PDAC—an indication where dominant escape axes shift from epitope-level mutation toward functional TME exclusion (risk score 85/100), antigen heterogeneity and clonal outgrowth (70/100), and soluble antigen sink (55/100). This risk stratification reshapes construct design logic: Discovery 4.0 recommends OR-gate dual-target designs (MSLN + CLDN18.2 or TROP2) to hedge against clonal heterogeneity, paired with biepitopic within-MSLN designs (E4+E6 or E5+E4) to address glycoform masking and soluble antigen competition. The Cell Suitability Engine shifts top recommendations toward iNKT cells (82/100) and iPSC-NK (74/100) for their superior TME infiltration capacity. Platform generalisation is achieved by substituting only antigen-specific inputs—isoform catalog, structure ensemble, mutation priors, shedding parameters—while the full computational stack transfers unchanged.

## Discussion

Discovery 4.0 establishes antigen escape resistance as a prospectively optimised engineering objective rather than a reactive clinical concern. Three mechanistic insights underlie its performance. First, epitope selection under mutation and selection pressure is a more powerful lever for escape resistance than affinity optimisation alone: selecting E4 over E1 for CD19 reduces CEL by approximately 65% before any multivalent architecture benefit is applied. Second, multivalent constructs reduce escape probability multiplicatively—a consequence of requiring simultaneous, independent loss of two or three distinct binder-epitope interactions, a combinatorial barrier that exponentially outpaces sequential resistance evolution. Third, biophysical compatibility between binders and escape-resistant epitopes is self-reinforcing: structurally indispensable epitopes require binders with kinetics—sub-nM KD, low Koff, low steric occlusion—that are intrinsically more developable.

The 98.1% escape reduction achieved by CAR_TRI_CD19E4·CD20E3·CD22E4 is significant in the context of B-cell malignancy, where sequential antigen-loss relapse after single-target and dual-target CAR therapy is increasingly the dominant failure mode [14, 15, 22]. By requiring tumour cells to simultaneously lose three independently escape-proofed interactions at epitopes that cannot be mutated without destabilising antigen folding, Discovery 4.0 raises the combinatorial barrier to escape beyond what is achievable through empirical therapy escalation.

In silico binder and construct predictions require full dose-response characterisation, in vivo pharmacokinetic validation, and long-term animal model studies before clinical translation.

Discovery 4.0’s self-improving architecture—in which experimental validation data are incorporated as training feedback to progressively refine the Mutation–Epitope Risk Engine and Cell Suitability scoring—positions the platform for continuous calibration as clinical evidence accumulates. Modular generalisation to new antigens requires substituting only antigen-specific inputs, enabling rapid deployment across oncological indications (solid tumours including PDAC, NSCLC, colorectal cancer) and autoimmune diseases where CAR-based tolerance induction is being explored.

In summary, Discovery 4.0 demonstrates that antigen escape—long treated as inevitable—can be substantially mitigated through prospective computational design. Significant escape reduction confirmed across ∼100 in vitro experiments, establishes a new performance benchmark and provides a translational foundation for next-generation clinical candidates at Pioneera Biosciences.

## Methods

### Protein/Epitope Foundation Encoder (Layer 1)

Antigen sequences for CD19, CD20, CD22, and BCMA were obtained from UniProt (P15391, P11836, P20273, Q02223) and supplemented with all deposited structural coordinates from the Protein Data Bank. Protein language model embeddings were generated using ESM-2 architecture variants, producing contextual residue representations capturing co-evolutionary information. Conformational epitope graphs were constructed by identifying surface-accessible patches (SASA > 20 Å^2^) and computing contact neighbourhoods within 8 Å of each patch centre. Three to eight candidate epitope patches were defined per antigen based on literature precedent and structural accessibility criteria.

### Mutation–Epitope Risk Engine (Layer 2)

Mutation priors were assembled from population-level variant databases (gnomAD v3.1, ClinVar) filtered for surface-exposed residues; therapy-induced selection composites derived from published relapse sequencing datasets (CD19-directed CAR-T, n=47 relapse samples; BCMA-directed therapies, n=28; CD20-directed, n=31); and in silico saturation mutagenesis using the Rosetta ddG protocol. A residue graph neural network was trained on this composite dataset to predict per-residue mutation frequency (%), structural destabilisation score (0–1), and immune selection pressure (0–1). Clinical Escape Likelihood (CEL; 0–1) was learned as a composite risk metric using gradient-boosted regression on held-out clinical escape events. Mutation Resistance Index (MRI; normalised 0–1 within each antigen) was computed as a weighted composite of conservation, structural anchoring, inverse mutation frequency, and inverse structural destabilisation.

### Virtual Binder Screening (Layer 3)

For each antigen, 5,000 synthetic binders were generated spanning scFv, VHH, and HCAb formats using a constrained sequence generation model trained on SAbDab and ABodyBuilder. Binding free energy and kinetic constants (Kon, Koff, KD) were predicted using a structure-based docking surrogate trained on SPR/ITC datasets. Off-target risk was assessed by sequence similarity to human proteins and structural complementarity screening against 150 off-target surface proteins. Composite Screen Score = 0.35 × normalised KD + 0.35 × mutation robustness + 0.20 × (1 − off-target risk) + 0.10 × manufacturability score.

### Multivalent Construct Design and Escape Modelling (Layer 4)

Construct generation explored tandem, parallel, and tri-specific configurations for all pairwise and triple combinations of top-ranked binder-epitope pairs. Avidity gain was predicted using a kinetic avidity model accounting for binder format, linker length, and spatial geometry of simultaneous engagement. Effective escape probability was computed as the product of per-epitope escape likelihoods under the assumption of independent escape events, with an empirical correction factor for spatial correlation within the same antigen. Escape Reduction vs. Best Mono = (1 − P_construct / P_best_mono) × 100%.

### Cell Suitability Engine (Layer 5)

Eight effector subtypes were modelled using phenotype-anchored parameter priors: CD4□ TCM, CD4□ TEM, CD8□ TCM, CD8□ TEMRA, NK CD56bright, NK CD56dim, iPSC-derived NK, and γδ T cells. For hematologic indications, composite suitability scores weighted persistence (25%), exhaustion resistance (25%), cytotoxic capacity (30%), and cytokine profile (20%). For solid tumour indications, weights shifted toward infiltration likelihood (20%), TME resistance (20%), with additional contributions from allogeneic feasibility, manufacturability, and safety controllability.

### In Vitro Co-incubation Validation

Antigen-expressing target cell lines (CD19□, CD20□, CD22□, BCMA□) were co-incubated with CAR-expressing primary T or NK cells at effector-to-target ratios of 1:1, 2:1, and 5:1. Escape was measured as the percentage of surviving target cells that lost surface antigen expression at 72 and 96 hours, quantified by flow cytometry with orthogonal antigen-specific antibodies. Escape Ratio = [% antigen-negative survivors (conventional CAR)] / [% antigen-negative survivors (Discovery 4.0 construct)] at matched conditions. Statistical comparisons used Mann– Whitney U tests with Bonferroni correction.

## Data Availability

Epitope scoring tables, binder screening outputs, construct design parameters, and in vitro co-incubation datasets are available from the corresponding author upon reasonable request subject to a material transfer agreement. The Discovery 4.0 platform is proprietary; key algorithmic parameters are described in Methods.

